# Computational and experimental analysis of the glycophosphatidylinositol-anchored proteome of the human parasitic nematode *Brugia malayi*

**DOI:** 10.1101/625111

**Authors:** Fana B. Mersha, Leslie K. Cortes, Ashley N. Luck, Colleen M. McClung, Cristian I. Ruse, Christopher H. Taron, Jeremy M. Foster

## Abstract

Further characterization of essential systems in the parasitic filarial nematode *Brugia malayi* is needed to better understand its biology, its interaction with its hosts, and to identify critical components that can be exploited to develop novel treatments. The production of glycophosphatidylinositol-anchored proteins (GPI-APs) is essential in humans, yeast, and the nematode *Caenorhabditis elegans*. In addition, GPI-APs perform many important roles for cells. In this study, we characterized the *B. malayi* GPI-anchored proteome using both computational and experimental approaches. We used bioinformatic strategies to show the presence or absence of *B. malayi* GPI-AP biosynthetic pathway genes and to compile a putative *B. malayi* GPI-AP proteome using available prediction programs. We verified these *in silico* analyses using proteomics to identify GPI-AP candidates prepared from the surface of intact worms and from membrane enriched extracts. Our study represents the first description of the GPI-anchored proteome in *B. malayi* and lays the groundwork for further exploration of this essential protein modification as a target for novel anthelmintic therapeutic strategies.

## Introduction

*Brugia malayi* is one of three related parasitic filarial nematodes responsible for human lymphatic filariasis, a disease that threatens more than 800 million people in 54 countries [1]. Adult worms reside in the lymphatic system where they can survive for an average of 6 to 8 years in immunocompetent individuals. Female worms release millions of progeny (microfilariae) during their lifetime which enter the bloodstream from where they can be ingested by feeding mosquitos. After larval molting and development within the insect vector, infective third stage larvae upon a blood meal can be transmitted to other humans in which they undergo further molting as they mature and migrate to the lymphatics thus completing the life cycle [2, 3]. Human lymphatic filariasis infections range from being asymptomatic to causing severe chronic symptoms that include damage to the lymphatic system and kidneys, and eventual clogging of lymphatic vessels with swelling of genitalia and limbs (elephantiasis) [2, 3]. Although more than 850 million people received treatments for lymphatic filariasis as part of the WHO effort to eliminate the disease [1], improvement of the current drug options would facilitate better control measures. Current treatments are only effective at reducing the levels of microfilariae circulating in blood so that further transmission is prevented, but they do not kill adult worms which cause the severe symptoms associated with this disease [1, 4]. In addition, there is evidence of emerging resistance to drugs used in current treatments of filarial infections [5, 6].

Many proteins are covalently attached to the surface of eukaryotic cells via a glycophosphatidylinositol (GPI) anchor. GPI precursors are synthesized in a step wise manner onto a phosphatidylinositol (PI) lipid moiety in the endoplasmic reticulum and transferred *en bloc* to proteins through the action of over 20 biosynthetic proteins. GPI-anchored proteins (GPI-APs) are transported via the secretory pathway and ultimately reside on the extracellular face of the plasma membrane and in lipid rafts [7]. This GPI-AP biosynthetic pathway is presumed to be ubiquitous in eukaryotes, and has been widely studied in mammals [8], fungi [9], plants[10] and protozoa [11, 12].

GPI anchors are critical for the proper localization and function of their appended proteins [13]. In mammals, interruption of GPI anchor biosynthesis during development is embryonic lethal [14] and an acquired defect in GPI anchoring causes the human hemolytic disease paroxysmal nocturnal hemoglobinuria (PNH) [8]. In lower eukaryotes, GPI anchoring is essential in the yeast *Saccharomyces cerevisiae* [15] and in the bloodstream form of the protozoan parasite *Trypanosoma brucei* [16]. Generation of gene knockout and conditional mutations the GPI-AP biosynthetic pathway genes in the free-living nematode *Caenorhabditis elegans* showed that GPI-anchoring is essential for egg and oocyte development [17], and for epithelial integrity during embryogenesis [17, 18]. Additionally, in the plant *Arabidopsis thaliana*, GPIs are critical for proper cell wall formation and coordinated cell growth [10]. The phylogenetic breadth of these studies points toward the process of GPI-anchoring of proteins being critical throughout eukaryotic cell biology.

Many types of plasma membrane proteins possess GPI anchors. Examples include adhesion molecules, surface receptors, prions, cell wall structural proteins and various types of enzymes [13]. In protozoan human parasites, GPI-APs are critical determinants of host-parasite interactions. For example, in *T. brucei,* the highly variable GPI-anchored variant surface glycoprotein coats the cell surface and allows the parasite to evade the host immune system [19]. In the malarial parasite *Plasmodium falciparum*, infectious merozoites are coated mainly with GPI-anchored MSP-1 and MSP-2 proteins [20]. Though MSP-2 function is poorly understood, MSP-1 plays a role in the binding and invasion of red blood cells by the merozoite. As such, MSP-1 has been identified as an antigen for naturally acquired immunity against malaria and is being investigated as a potential vaccine candidate [20, 21]. These studies highlight the importance of understanding the repertoire of GPI-APs present in parasites. However, little is known about the GPI assembly and the GPI-anchored proteome in filarial parasitic nematodes such as *B. malayi*.

Here, we provide the first characterization of the GPI-anchored proteome of *B. malayi*. We report the conservation of GPI biosynthesis pathway genes and the prediction of putative GPI-anchored proteins encoded in the *B. malayi* genome. We further confirm the presence of *B. malayi* proteins that have the GPI anchor post-translational modification and identify *B. malayi* GPI-AP using enrichment techniques and proteomic analysis. Our study gives a better understanding of the *B. malayi* GPI-anchored proteome and the roles for GPI-APs and may ultimately help identify new therapeutic or diagnostic targets.

## Materials & methods

### GPI-AP synthesis pathway prediction

Human and *C. elegans* proteins encoded by each gene in the GPI-AP synthesis pathway [17] were used to search for orthologs in the *B. malayi* genome at Wormbase (release WS265) using BLASTP and TBLASTN (https://www.wormbase.org/tools/blast_blat)[22] and also cross checked against the KEGG GPI-AP synthesis pathway [23] and against Pfam [24]. For the three orthologs not identified using human and *C. elegans* proteins, Pfam domains of PIG-H (PF10181), PIG-Y (PF15159), and DPM2 (PF07297) were used to find the top ten most diverse members of the family identified on NCBI Conserved Protein Domain Database [25] and those proteins were used to search for orthologs in the *C. elegans* and *B. malayi* genomes both using BLASTP at NCBI [26] and Wormbase. Similarly, a taxonomic neighbor with an identified PIG-H (Aedes aegypti Q16JH1), PIG-Y (*Anopheles minimus* A0A182WPC2) and DPM2 (*Steinernema glaseri* A0A1I7ZJZ8) was used to search for orthologs. No matches for these three missing orthologs were found.

### FLAER Dot blot

FLAER (Alexa 488 proaerolysin variant) was obtained from Cedarlane Labs (Burlington, Canada). *Escherichia coli* strain OP50 was cultured overnight in liquid LB media. *C. elegans* N2 Bristol was cultured as previously described [27]. Total lysates for *E. coli* OP50 and *C. elegans* were prepared by sonication in cold sucrose buffer (10 mM Tris-HCl, pH 7.5, 250 mM sucrose, 1 mM EDTA) with 1X Calbiochem protease inhibitor cocktail set III (MilliporeSigma, Burlington MA). Soluble lysates were prepared by collecting the supernatant after centrifugation of the lysate at 1000 x g for 10 minutes at 4°C. *B. malayi* adult worms (TRS Labs Inc., Athens GA) were cultured in plates containing 40 ml GIBCO RPMI-1640 culture medium (ThermoFisher Scientific, Waltham MA) supplemented with 0.4 ml of 100X Antibiotic Antimycotic Solution which contains 10,000 units penicillin, 10 mg streptomycin and 25 μg amphotericin B per ml (Sigma, St. Louis, MO, USA). Each plate had approximately 100 female worms and 90 male worms. After 48 hours at 37°C in 5% CO_2_ in air, worms were washed 4 times in 10 ml of the same medium. Worms from two plates were transferred to new plates with 5 ml medium and served as the mock control while worms from a different set of two plates were transferred to new plates with 5 ml culture medium containing 15 μg *Bacillus cereus* phosphatidylinositol-specific phospholipase C (PI-PLC) [28]. After incubation for 30 minutes at 37°C, the medium was collected and worms were washed with an additional 10 ml fresh medium. The samples containing the released proteins for either mock treated or PI-PLC treated were pooled and used to prepare *B. malayi* surface fractions for LC-MS/MS analysis described below. To prepare *B. malayi* total lysate, the washed worms were resuspended in 2 ml cold sucrose buffer with 1X Calbiochem protease inhibitor cocktail set III (MilliporeSigma) and homogenized using Precellys SK38 beads (Bertin Instruments, Rockville, MD) in a MiniLys homogenizer (Bertin Instruments, Rockville, MD) by agitating at 5000 rpm for 10 seconds eight times and cooling samples on ice between each homogenization. Debris was pelleted at 1000 x g, 4°C, 10 minutes and the total lysate transferred to a new tube. To ensure complete lysis and to wash the beads, homogenization was repeated on pellet/debris and like samples were pooled. The protein concentration was measured by Bradford assay (Bio-Rad, Hercules, CA) and samples were spotted in a 1:1 dilution series on nitrocellulose membrane. The *C. elegans* lysates served as a positive control while *E. coli* lysates, bovine serum albumin (New England Biolabs) and buffer served as negative controls. After the samples dried, the nitrocellulose membrane was washed in binding buffer (50 mM NaH_2_PO_4_, pH 7.5, and 0.3% TWEEN 20) for 20 minutes and incubated overnight with 3 nM FLAER in the same buffer at 4°C. The membrane was washed six times in binding buffer and fluorescence detected on an Amersham Typhoon imager (GE Healthcare Life Sciences).

### Predicted *B. malayi* GPI-APome

The *B. malayi* proteome (brugia_malayi.PRJNA10729.WBPS10.protein.fa) was downloaded from Wormbase in May 2018 and analyzed for GPI-APs with three GPI-AP prediction tools:

GPI-SOM (http://gpi.unibe.ch)[29]

PredGPI (http://gpcr.biocomp.unibo.it/predgpi/pred.htm)[30]

big-PI (http://mendel.imp.ac.at/gpi/gpi_server.html)[31].

For PREDGPI and big-PI, we further restricted the data set by using Signal-P 4.1 (http://www.cbs.dtu.dk/services/SignalP/)[32] to require that the proteins have a N-terminal signal sequence. For all three datasets, we also ran the transmembrane prediction program TMHMM v2.0 (http://www.cbs.dtu.dk/services/TMHMM) [33] and removed any proteins that had more than three predicted transmembrane domains to eliminate multipass transmembrane proteins.

To compare the predicted GPI-APs to previously published proteomic studies which use pub_locus ID, we cross-referenced the Wormbase and Uniprot databases to correlate Wormbase ID to the PUB_loci/pub_locus numbers.

### Protein sample preparation for LC-MS/MS

#### *B. malayi* surface fractions

The pooled supernatant and wash from the mock treated worms and the PI-PLC treated worms described above to prepare *B. malayi* samples for FLAER Dot Blot were collected. The pools were each filtered through a 0.22-micron filter and concentrated using VivaSpin 10,000 MWCO concentrator (GE Healthcare Life Sciences).

#### *B. malayi* membrane fractions

150 *B. malayi* adult female worms were resuspended in 2 ml cold sucrose buffer with 1X Calbiochem protease inhibitor cocktail set III (MilliporeSigma), homogenized, and total lysate was prepared as described above. The lysate was then centrifuged at 100,000 x g for 1 hour at 4°C to pellet the membrane fraction away from cytosolic supernatant [34]. The resulting pellet was washed with 10 ml of 150 mM Tris-HCl, pH 8.0 and ultracentrifugation was repeated. The pellet containing the washed membrane fraction was collected. The washed membrane pellet was resuspended in 2 ml cold 1X phosphate buffered saline and divided into 2 x 1ml aliquots. 3 μg PI-PLC [28] was added to one aliquot and both samples were incubated at 37°C for 30 minutes. An equal volume of sucrose buffer was added to each sample to fill the ultracentrifuge tubes and both samples were centrifuged at 100,000 x g for 1 hour at 4°C. The supernatants were collected and concentrated using VivaSpin 5000 MWCO (GE Healthcare) concentrators and then dialyzed into 25 mM ammonium bicarbonate, pH 7.5 containing 40 mM NaCl. An additional washed membrane sample was prepared from 18 female and 12 male worms as described above without the PI-PLC treatment and then GPI-AP enriched as previously described [35]. Briefly, the washed membrane pellet was resuspended in 0.1 ml water and then brought to 10:10:3 dichloromethane:methanol: water by adding 0.7 ml 1:1 dichloromethane:methanol until the pellet was solubilized. Then 1:1 methanol:water in 0.1 ml increments was added until a phase separation occurred. The sample was centrifuged at 2000 x g for 10 minutes. The upper aqueous phase and interphase was collected and the extraction was repeated by adding one volume dichloromethane. The upper aqueous phase was collected and dried to a pellet in a Speed Vac. This delipidated pellet was resuspended in 0.1 ml PBS and mixed with an equal volume of water-saturated butanol and centrifuged at 1000 x g for 10 minutes after mixing well. The upper butanol phase was removed and the extraction was repeated on the lower aqueous phase. The aqueous phase containing the GPI-AP enriched sample from the second butanol extraction was collected and dried in a Speed Vac.

### Mass Spectrometry and Analysis

#### Surface samples

Surface samples were digested with Trypsin-ultra, Mass Spectrometry Grade (New England Biolabs, Ipswich MA) overnight at 37°C using a filter aided sample preparation kit (Expedeon, San Diego CA). For a complete protocol see [36]. Each sample was analyzed using an automated 3-step MudPIT protocol on a Q Exactive mass spectrometer (ThermoFisher Scientific, Waltham MA) using HCD fragmentation.

#### Membrane samples

Membrane samples were digested with Trypsin-ultra, Mass Spectrometry Grade (New England Biolabs) upon addition of ProteaseMax (Promega, Madison WI) for 3 hours at 37°C. Each sample was analyzed using an 8-step MudPIT protocol on an LTQ-Orbitrap XL (ThermoFisher Scientific) using CID fragmentation.

#### LC-MS/MS Data Analysis

Spectral data were searched against the combined *B. malayi* proteome (brugia_malayi.PRJNA10729.WBPS10.protein.fa) from Wormbase [22] and the *Wolbachia* endosymbiont of *B. malayi* proteome from NCBI [26], both downloaded in May 2018 and then analyzed using Byonic software (Protein Metrics, Cupertino, CA). Common contaminants such as keratins, caseins, trypsin, and BSA were removed from the analysis. Protein output was set at 1% FDR. Other parameters used for each Mass Spectrometry dataset:

QExactiveQTOF: Cleavage site= RK; Cterminal side=Fully specific; missed cleavage=2; Mass tolerance=10 ppm; QTOF/HCD fragmentation with 20 ppm fragment mass tolerance. Fixed modifications=carbamidomethyl @ C/+57.021464. Variable Modifications=Oxidation/ +15.994915 @ M, Deamidation /+0.984016 @ N,Q, Carbamyl @ K/ 43.005814.

LTQ-Orbitrap XL: Cleavage site= RK; Cterminal side=Fully specific; missed cleavage=2; Mass tolerance=20 ppm; CID low energy fragmentation with 0.6 Dalton fragment mass tolerance. Fixed modifications=carbamidomethyl @ C/+57.021464. Oxidation/+15.994915 @ M, Deamidation /+0.984016 @ N,Q.

All protein matches that were limited to one unique peptide are highlighted in red in S2-S6 Tables and were not used in further protein identification analysis.

To compare the LC-MS/MS proteins to published proteomic studies that used different protein IDs, we cross referenced the Wormbase [22] and UniProt [37] databases to correlate Wormbase ID from our data to the PUB_loci/pub_locus numbers or to UniProtKB ID for the *B. malayi* proteome and to the UniProt database to correlate NCBI ID with UniProtKB ID or wBm gene numbers for the *Wolbachia* proteome. The synonyms we found associated with each protein in our dataset are listed in S8 Table.

## Results and discussion

### *B. malayi* GPI-AP biosynthesis

While the overall process for forming GPIs and attaching them to proteins is remarkably well-conserved across eukaryotes, organism-specific differences are known and have been previously reviewed [38]. Thus, it is possible to gain initial insights into the various steps of GPI anchoring through examination of a genome for the presence or absence of GPI synthesis machinery. Using protein sequence homology, the GPI synthesis pathway of *C. elegans* was predicted [17]. Here, we conducted a similar analysis to identify *B. malayi* orthologs and additional *C. elegans* orthologs of the GPI-AP synthesis pathway genes encoded in the their respective genomes using the KEGG pathway [23], Pfam [24], and BLASTP [39] (Fig 1, Table 1).

**Fig 1.**
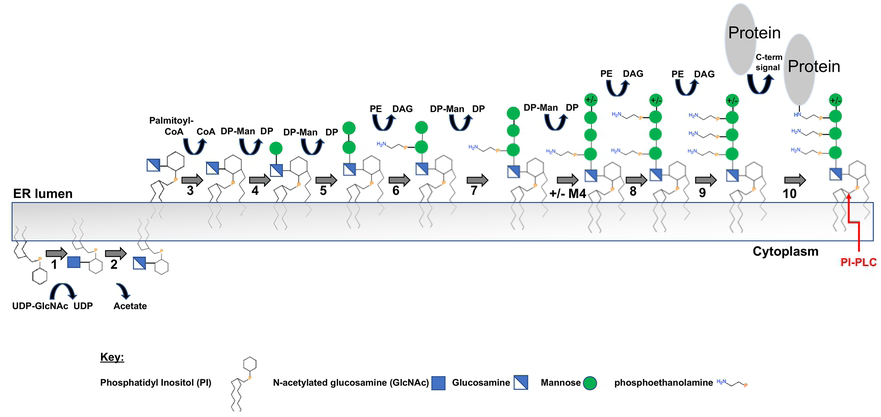
The conserved steps in the GPI-AP synthesis pathway. The enzymes or enzyme complexes for each step are listed in Table 1. UDP-GlcNAc: UDP-N-acetylglucosamine; DP-Man: Dolichol-P-Mannose; PE: Phosphatidylethanolamine; DAG: diacylglycerol. The phosphatidylinositol-specific phospholipase C (PI-PLC) cleavage site to release glycosylated protein is noted on the final structure.

**Table 1.** GPI-AP synthesis pathway.

|  | Step | Enzyme or Enzyme Complex | Human | *C. elegans* | *B. malayi* |
| --- | --- | --- | --- | --- | --- |
| GPI anchor synthesis and protein attachment | 1 | GPI-GlcNAc transferase | PIG-A | piga-1 (D2085.6) | Bm3057 |
|  |  |  | PIG-C | T20D3.8 | Bm5863 |
|  |  |  | PIG-H |  |  |
|  |  |  | PIG-P | Y48E1B.2 | Bm6655 |
|  |  |  | PIG-Q | F01G4.5 | Bm3139 |
|  |  |  | PIG-Y |  |  |
|  |  |  | DPM2 |  |  |
|  | 2 | GlcNAc-PI de-N-acetylase | PIG-L | Y52B11C.1 | Bm6741 |
|  | 3 | Inositol acyltransferase | PIG-W | Y110A2AL.12 | Bm7272 |
| | 4 | $\alpha$ 1-4 mannosyltransferase I | PIG-M | B0491.1 | Bm1821 |
|  |  |  | PIG-X | F49E7.2 | Bm4227 |
| | 5 | $\alpha$ 1-6 mannosyltransferase II | PIG-V | T09B4.1 | Bm5692 |
|  | 6 | EtNP transferase I to Man-1 | PIG-N | Y54E10BR.1 | Bm6800 |
| | 7 | $\alpha$ 1-2 mannosyltransferase III | PIG-B | T27F7.4 | Bm13907 |
| | +/- M4 | $\alpha$ 1-2 mannosyltransferase IV | PIG-Z | ZC513.5 | Bm7406 |
|  | 8 | EtNP transferase III to Man-3 | PIG-O | C27A12.9 | Bm2424 |
|  |  |  | PIG-F | F49E8.1c | Bm18036 |
|  | 9 | EtNP transferase II to Man-2 | PIG-G | F28C6.4 | Bm3669 |
|  |  |  | PIG-F | F49E8.1c | Bm18036 |
|  | 10 | GPI transamidase | PIG-K | T05E11.6 | Bm13834 |
|  |  |  | GAA1 | F33D11.9 | Bm3795 |
|  |  |  | PIG-S | T14G10.7 | Bm5798 |
|  |  |  | PIG-T | F17C11.7 | Bm3419 |
|  |  |  | PIG-U | T22C1.3 | Bm5915 |
| Remodeling | 11 | GPI deacylase | PGAP1 | T19B10.8 | Bm5838 |
|  | 12 | Metallophosphoesterase 1 | PGAP5 | B0511.13 | Bm1843 |
|  | 13 | GPI-specific phospholipase A2 | PGAP3 | R01B10.4 | Bm5161 |
|  | 14 | GPI post attachment protein 2 | PGAP2 | tag-189 (T04A8.12) | Bm5533 |
| Mannose Donor |  | Dolichol-phosphate-mannosyltransferase | DPM1 | dpm-1 (Y66H1A.2) | Bm6987 |
|  |  |  | DPM2 |  |  |
|  |  |  | DPM3 | dpm-3 (F28D1.11) | Bm3675 |

Orthologs of human genes identified for the stepwise construction of GPI-APs as shown in Fig 1 are listed both for *C. elegans* and for *B. malayi* under the GPI anchor synthesis section (Steps 1-10). GlcNAc is N-acetylglucosamine; PI is phosphatidylinositol; EtNP is ethanolamine phosphate; Man is Mannose. Missing orthologs are shaded gray. Post-GPI attachment to proteins (PGAP) gene orthologs are listed in the remodeling section (Steps 11-14). Orthologs to genes involved in synthesis of the mannose donor, Dolichol-phosphate mannose are listed in the mannose donor section and described in S1 Fig.

Remarkably, orthologs to three genes encoding components of the first step, PIG-H, PIG-Y, and DPM2 are not found in either *B. malayi* or *C. elegans* genomes. These genes are part of the seven-member GPI-GlcNAc transferase complex that transfers the N-acetylated glucosamine (GlcNAc) to the lipid phosphatidylinositol (PI) on the cytoplasmic side of the endoplasmic reticulum (Step 1 in Fig 1 and Table 1). While DPM2’s role for stabilizing the complex is not essential, both PIG-H and PIG-Y directly interact with the catalytic subunit PIG-A and are critical for the activity of the complex [40]. Though the smaller size (PIG-H is 188 amino acids; PIG-Y is 71 amino acids; DPM2 is 84 amino acids) and relative hydrophobicity of these membrane proteins can make it more difficult to identify them by sequence homology, the absence of PIG-H and PIG-Y in all sequenced nematode genomes suggests that this represents a true difference between nematodes and other species. If the functions of these genes are carried out by analogous but divergent proteins, the filarial parasite GPI-GlcNAc transferase complex could differ enough from the human enzyme complex to be explored further as a therapeutic target.

As well as playing a stabilizing role in the GPI-GlcNAc transferase complex, DPM2 is a component of the dolichol phosphate mannosyltransferase (DPM) complex (Table 1). The mannose donor, dolichol-P-mannose (DP-Man in Fig 1) required for GPI anchor synthesis (Steps 4, 5, 7, 8 in Fig 1 and Table 1) as well as other glycosylation pathways, is synthesized by this enzyme complex [41, 42]. In humans, there are three proteins in this complex: DPM1, DPM2 and DPM3. DPM1 is the catalytic subunit bound to the membrane by DPM3 while DPM2 stabilizes DPM3 [41]. In *S. cerevisiae* and *T. brucei*, the complex is missing DPM3 but their respective DPM1 orthologs have a transmembrane domain which the human DPM1 does not have [42]. In contrast, *C. elegans* and *B. malayi* have a human-like DPM1 ortholog without a transmembrane domain and a DPM3 ortholog to bind the DPM1 to the membrane but no DPM2 ortholog was identified (S1 Fig).

*B. malayi* orthologs were identified for twenty one of twenty-four GPI anchor synthesis proteins and all four remodeling proteins. Excluding the three missing orthologs for the GPI-GlcNAc transferase complex, the pathway appears similar to the human and yeast GPI anchor synthetic pathways [9]. In addition to the mannosyltransferases that add the first (Man-1), second (Man-2), and third (Man-3) GPI core mannoses, a fourth mannosyltransferase (mannosyltransferase IV) was also identified in both *B. malayi* and *C. elegans*. Fourth mannose (Man-4) addition is essential in yeast, and although less common, Man-4 is present on some human GPI-APs but is absent from GPIs of *T. brucei* [43, 44]. Ethanolamine phosphate (EtN-P) transferases I, II and III, that add side-branching EtN-P moieties to Man-1, Man-2 and Man-3 of the GPI glycan, respectively, are each present in *B. malayi*. EtN-P transferases I and II are found in yeast and mammals but are absent in Trypanosomes [38]. A carboxy-terminal GPI attachment signal sequence directs the transfer of the GPI precursor to a protein’s omega site via the GPI-transamidase enzyme complex. The result is a covalent amide linkage between the protein and the phosphoethanolamine moiety of the GPI anchor [11]. Thus, equivalent to the human and *C. elegans* pathway, the *B. malayi* GPI anchor synthesis pathway has the enzymes needed to build and transfer to proteins a trimannosyl- (and possibly tetramannosyl-) GPI precursor bearing EtN-P on its innermost two mannoses (Fig 1). Additionally, *B. malayi* orthologs were also identified for Post GPI attachment proteins (PGAP) that can remodel the GPI anchor after it is attached to the protein.

### Detection of GPI-APs in *B. malayi*

To verify that GPI-APs are present in *B. malayi*, we used FLAER, an Alexa 488-labeled proaerolysin variant, that binds specifically to GPI-APs [45] and in *C. elegans* showed the presence of GPI-APs in germline cells and somatic cells [17]. In a dot blot, the total extracts from both *C. elegans* and *B. malayi* adult worms were bound by FLAER and produced a signal that was dependent on protein concentration indicating the presence of GPI-APs in these samples (Fig 2). *E. coli* extracts, purified BSA and buffer controls did not react with FLAER showing only background signal. A dot blot of the same samples with no FLAER showed no autofluorescence (S2 Fig). This data demonstrates that GPI-APs are present in *B. malayi* and that the structure of the GPI sugar core is sufficiently similar to previously studied GPI-APs at least in the glycophosphatidylinositol moiety as aerolysin requires both polypeptide and GPI, but not lipid, for binding [46].

**Fig 2.**
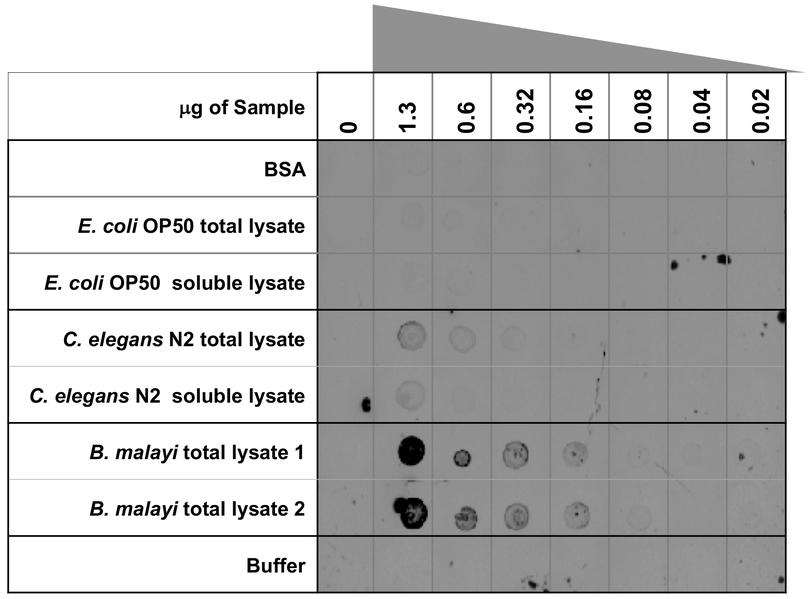
FLAER Dot Blot. *B. malayi* and *C. elegans* lysates react with FLAER in a concentration dependent manner. Samples: BSA, *E.coli OP50* total and soluble lysate, *C. elegans* N2 total and soluble lysate, *B. malayi* total lysate 1: *B. malayi* total lysate untreated, *B. malayi* total lysate 2: *B. malayi* surface PI-PLC treated total lysate. After 1:1 series dilution, samples were spotted on to a nitrocellulose membrane and incubated overnight with FLAER reagent.

The FLAER signal for both *B. malayi* lysates was stronger than those of *C. elegans* even though all the samples were normalized by protein concentration. While it is possible that *B. malayi* produce more GPI-APs than free living nematodes, this observation could also be due to a difference in reproductive production. Female *B. malayi* are estimated to release tens of thousands of microfilaria per day [47, 48] whereas *C. elegans* release a maximum of 1000 progeny during their 3 day lifecycle [49]. The greater number of eggs, embryos and/or immature microfilariae present in *B. malayi* adult females may be responsible for the higher level of GPI-AP signal as it was shown in *C. elegans* that germline cells stained brightly with FLAER [17]. Finally, there was no discernable decrease in signal when comparing total extracts prepared from *B. malayi* that were untreated (*B. malayi* total lysate 1) to those that were treated with phosphatidylinositol-specific phospholipase C (PI-PLC) (*B. malayi* total lysate 2) prior to homogenization and lysis. It is possible that cleavage of the surface GPI-APs by PI-PLC was not efficient or was inhibited due to limited access of PI-PLC to GPI-AP either by the surface coat of *B. malayi* or clustering of GPI-AP in lipid rafts. Also, GPI-APs on the adult *B. malayi* surface might represent only a small proportion of the total pool of GPI-APs which includes not only the all the GPI-AP present in non-surface cells but also all the immature progeny as well. It is estimated that *B. malayi* have approximately 2000 somatic cells but the majority of the signal in the total extract of female adult *B. malayi* may derive from the tens of thousands of eggs, embryos and/or immature microfilariae [48]. Lastly, normal turnover or endogenous PI-PLC may have already released most GPI-APs on the worm surface. Nevertheless, the *B. malayi* total extracts did react with FLAER demonstrating the presence of GPI-AP in these filarial nematodes.

### Computational prediction of the *B. malayi* GPI-APome

The *B. malayi* GPI-anchored proteome (GPI-APome) was assembled using available GPI-AP prediction algorithms. Three programs (GPI-SOM, PredGPI, and bigPI) were used to search the *B. malayi* proteome and identify 210 predicted GPI-APs (S1 Table). The data from each program is plotted in Fig 3A using UpSetR analysis [50] to compare the intersecting sets. Sixteen proteins were identified by all three programs; two or more programs identified 62 proteins; 148 proteins were identified by a single prediction program. These 210 predicted GPI-APs represent 1.5% of the 13,409 proteins in the *B. malayi* predicted proteome. This is comparable to other computationally predicted GPI anchored proteomes [30, 51, 52].

**Fig 3.**
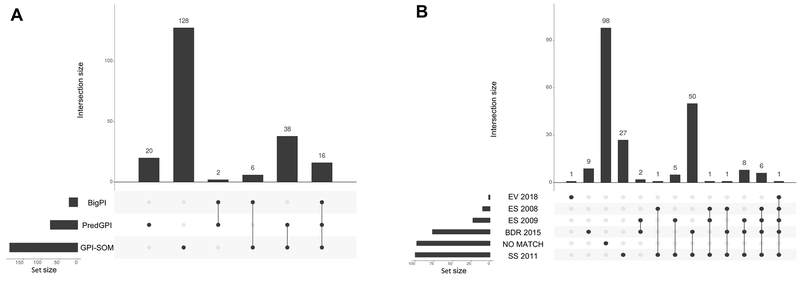
UpSetR analysis of *B. malayi* predicted GPI-APome and proteomic comparison. **(A) Predicted GPI-APs from GPI-SOM, PredGPI and bigPI.** Over half of the proteins are exclusive to either PredGPI or GPI-SOM prediction sets and are designated with a single dot. 16 proteins were identified by all three GPI-AP prediction programs as indicated by three linked black circles and 46 were identified by two GPI-AP prediction programs as indicated by two linked black circles. **(B) The predicted GPI-APome was compared to previous *B. malayi* proteomic datasets.** EV 2018 [53] is an extracellular vesicle proteome with one predicted GPI-AP in common with all other proteomes and one predicted GPI-AP exclusive to itself. ES 2008 [54] and ES 2009 [55] are the excretory and secretory proteomes and BDR 2015 [56] is the body wall, the digestive tract and the reproductive tract proteomes. SS 2011 [57] is the stage specific proteome set. No Match indicates GPI-APome proteins not found in these five proteomic studies. The set size shown on the left of each graph indicates the number of proteins in each set.

Of the 210 predicted GPI-APs, 112 were detected in previous *B. malayi* proteomic studies (Fig 3B, S1 Table). Protein identifications are those reported by each individual study. All of the predicted GPI-AP matches except for one, were found in the body wall, digestive, and reproductive proteomic analysis [56] and/or the stage specific proteome study [57]. Only 25 predicted GPI-APs were found in excretory or secretory proteomes [54, 55] with no predicted GPI-APs exclusive to these proteomes. Two predicted GPI-APs were found in the recent adult extracellular vesicle proteome [53] with one protein exclusive to that proteome. Thus, roughly half of our computationally predicted GPI-APome comprises proteins that have been found in proteomes expected to harbor GPI-APs.

Conversely, 98 of the predicted *B. malayi* GPI-APs, noted as “No Match*”* in Fig 3B, have not been identified in any previous *B. malayi* proteomic study. Forty-six of these proteins have synonym IDs allowing for comparison to previous proteomes but they were not found in those studies. Possibly, some GPI-APs are only expressed transiently or at life stages not investigated in earlier studies or that protein abundance for those predicted GPI-AP was not high enough for identification. However, the remaining 52 proteins did not have a synonym ID and are likely from newly defined genes or gene models with updated genome versions since the previous proteomes were analyzed.

Bm7382, microfilaria surface antigen (other synonyms: Bm-103, Bm1_01550, Bma-MSA-1), was predicted to be a GPI-AP by two programs (S1 Table) and has been shown to localize to the hypodermis and cuticle of *B. malayi* adult female worms, L3 larvae, and microfiliariae [58]. This protein has already been in a vaccine trial utilizing the Mongolian gerbil model for *B. malayi* individually and as a fusion with another protein with promising results [58]. This further supports the idea that GPI-APs may make promising therapeutic targets.

### Mass spectrometry identification of potential *B. malayi* GPI-APs

A proteomics strategy was used to identify putative GPI-APs from adult *B. malayi*. Three different sample types were prepared for analysis. Firstly, the surface of intact adult worms was treated with PI-PLC to enzymatically release and solubilize of the protein away from the lipid moiety [59]. (See Fig 1 for cleavage site position). A mock-treatment with no PI-PLC was performed as a negative control. Secondly, a membrane fraction of *B. malayi* adult female worms was prepared by ultracentrifugation of a total lysate in a sucrose buffer to separate membrane proteins from soluble proteins. This membrane fraction was also treated with PI-PLC or mock-treated without PI-PLC as a negative control. Lastly, a GPI-AP enriched sample was prepared by performing a series of organic solvent partitions [35] to extract GPI-APs from a membrane fraction. This sample was not treated with PI-PLC. Proteins in all three samples types were digested with trypsin and the resulting peptides analyzed by LC-MS/MS.

The data from analysis of these five samples was used to identify peptides corresponding to ∼ 1400 proteins in the predicted *B. malayi* proteome and 27 from its endosymbiont *Wolbachia* proteome (S2 to S6 Tables). By excluding all proteins having a single unique peptide and reducing isoforms to one protein, a reduced dataset of 1012 proteins was compiled (S7 Table). We used UpSetR [50] to visualize the intersections of these identified proteins in our five samples (Fig 4). The membrane samples yielded the highest number of identified proteins with a total of 890 proteins; 653 proteins were identified in the membrane mock control sample; 680 were identified in the membrane PI-PLC treated sample; 101 proteins were identified in the organic solvent GPI-AP enriched sample (“Membrane Enrich”). The parasite surface samples yielded a total of 333 proteins with 281 proteins identified in the surface mock control sample and 209 proteins identified in the PI-PLC treated surface sample.

**Fig 4.**
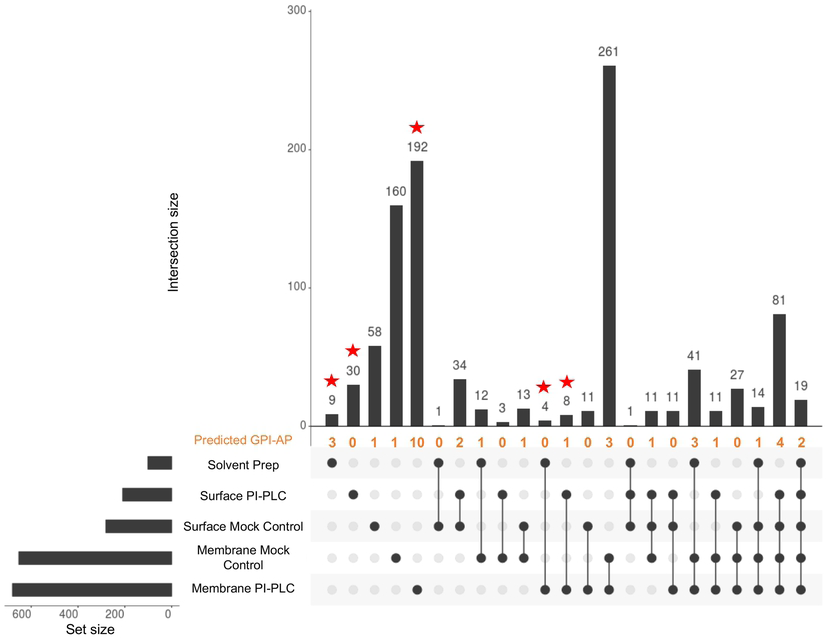
UpSetR data analysis of LC-MS/MS data from the five protein samples. **Membrane Enrich:** sample from membrane preparation that was enriched for GPI-APs using successive organic solvent partitioning; **Surface PI-PLC** and **Surface Mock Control**: released proteins from intact worms treated with PI-PLC or no enzyme; **Membrane Mock Control** and **Membrane PI-PLC:** released proteins from membrane preparation treated with no enzyme or PI-PLC. Starred columns represent sets only containing proteins from experimentally GPI-AP enriched samples. The number of predicted GPI-APs found in each dataset is indicated in orange.

The high number of proteins identified in the mock-treated control samples likely reflects the release of proteins and peptides through normal turnover processes, *B. malayi* proteases that remain in the sample, and possibly an endogenous PI-PLC (Bm5477). We had expected the PI-PLC treated samples to identify similar proteins as the mock controls and to identify additional GPI-APs that had been specifically released by the added enzyme. However, the small amount of protein recovered from intact worm surfaces, either with or without PI-PLC treatment, limited the depth of our LC-MS/MS analysis. Furthermore, it is possible that GPI-APs on intact worms are protected from PI-PLC cleavage due to their inclusion in densely packed lipid rafts or due to the surface coat.

Treatment of membrane fractions by PI-PLC produced different results. The membrane PI-PLC treated sample identified more proteins than its corresponding mock control sample (Fig 4, S5 and S6 Tables). The sets denoted by a red star in Fig 4 include proteins exclusively found in samples that should be enriched for GPI-APs (e.g., membrane PI-PLC, surface PI-PLC, and organic solvent GPI-AP enriched samples). The 243 proteins in these samples comprise experimentally enriched putative GPI-APs present in adult *B. malayi*.

To support the identification of GPI-APs, we compared the 1012 proteins from our LC-MS/MS data to the 210 proteins in our computationally predicted GPI-APome. Thirty-five candidate GPI-APs were present in both analyses and are listed in Table 2 and are noted in Fig 4 in orange as predicted GPI-AP. The membrane set had the highest number of GPI-APs exclusively found in PI-PLC treated or organic solvent-enriched GPI-AP samples (Table 2; highlighted boxes in the unique peptides column). This is also shown in Fig 4 where the membrane PI-PLC set contained more than 25% of computationally predicted GPI-APs while the remaining predicted GPI-APs were scattered throughout other sets. The surface PI-PLC exclusive set in Fig 4 showed no matches with the predicted GPI-APome, further supporting our conclusion that PI-PLC does not efficiently release GPI-APs from the surface of intact worms.

**Table 2.**
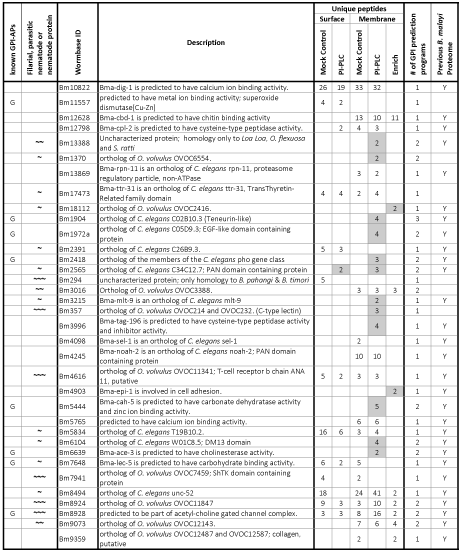
Correlation of computational and experimental identification of candidate *B. malayi* GPI-APs.

Proteins that are orthologs to previously identified GPI-APs are indicated by G in Column 1. Proteins that had greater than 50% BLASTP identity only to other nematodes, parasitic nematodes or parasitic filarial nematodes are designated respectively with ∼, ∼∼, ∼∼∼ in Column 2. The number of unique peptides identified in the five samples are listed in Columns 5-9 and are shaded if the identified protein was only present in PI-PLC or Enrich sample. The number of GPI-AP prediction programs that recognized each protein is listed in Column 10. Proteins found in previously published proteomes are designated with Y in the last column.

We searched for examples of previously identified GPI-APs amongst these thirty-five proteins to help validate our results. About 25% of the proteins are orthologs of proteins previously reported to be GPI-anchored in other organisms and are noted in Table 2. Phosphatase was the first GPI-AP to be identified by PI-PLC release [60]. Bm2418, a putative phosphatase, is predicted to be a GPI-AP by both GPI-SOM and PredGPI and is present in the *B. malayi* membrane PI-PLC sample. Bm2418 is an ortholog of *C. elegans* PHO-1 and PHO-4 proteins which were also identified as a GPI-APs in the *C. elegans* lipid raft proteome [61] and the *C. elegans* GPI-AP study [17], respectively.

Another well studied GPI-AP is acetylcholinesterase [62–64]. This enzyme activity has been previously characterized in *B. malayi* [65]. Here, we show that Bm6639 (Bma-ace-3), which is predicted to have acetylcholinesterase activity, is also predicted to be a GPI-AP by two programs and is present in the membrane PI-PLC sample. Bm11557, a third GPI-AP ortholog example, is a putative superoxide dismutase and was only found in the two surface samples. Superoxide dismutase has been shown to be a GPI-AP in the slime mold *Dictyostelium* where its role in chemotaxis was elucidated and because of its localization to the plasma membrane is thought to play a role in the concentration of extracellular superoxide and possibly in the flux of superoxide radicals into the cell [66]. Bm8928, predicted to be part of an acetyl-choline gated channel complex, is present in all five samples we analyzed by LC-MS/MS. It is predicted to be a GPI-AP in *B. malayi* by two programs and its *C. elegans* ortholog Y12A6A.1 (34% ID with Bm8928) was identified as a GPI-AP in the *C. elegans* lipid raft proteome (21). Bm7648 is another matching ortholog to a GPI-AP found in the *C. elegans* lipid raft proteome [61] and Bm1904, Bm1972a, Bm5444 are matching orthologs of GPI-APs identified in the *C. elegans* GPI-AP study [17]. These previously studied GPI-AP orthologs support the identity of the thirty-five *B. malayi* GPI-AP proteins we identified by the combination of both computational and experimental methods as being GPI-APs.

GPI-APs with potential diagnostic, therapeutic or vaccine applications given that they either are not present in the human proteome or are highly divergent are noted with **∼, ∼∼,** or **∼∼∼,** in Table 2. Bm294, Bm357, Bm4616, Bm7941, Bm8924, and Bm8928 have >50% identity only to proteins from other filarial nematodes (**∼∼∼**) like other *Brugia* spp. and *Onchocerca spp*. Bm13388, Bm3016, and Bm9073 have >50% identity only to proteins from other parasitic nematodes (**∼∼**) which include *Strongyloides* as well as filarial nematodes. Bm1370, Bm17473, Bm18112, Bm2391, Bm2565, Bm3215, Bm5834, Bm6104, Bm7648, and Bm8494 have >50% identity only to proteins found in nematodes (**∼**) including *C. elegans* but still remain distinct from proteins of other phyla. While the majority of these proteins remain uncharacterized, some information on their possible role is available, especially for the nematode-specific proteins with *C. elegans* orthologs. Bm3215 is an ortholog of *C. elegans* Mlt-9, a protein that has an important role in molting and has no human homolog. Mlt-9 is present in the hypodermis and seam at a critical time in worm development and RNAi silencing of mlt-9 disrupts L3 to L4 and L4 to adult molting [67]. Bm2565 is an ortholog of *C. elegans* C34C12.7, an uncharacterized protein that contains a PAN domain, which confers protein-protein or protein-carbohydrate interaction [68]. As well as finding Bm2565 in the membrane PI-PLC sample, this protein was also one of the few GPI-APs identified in the surface PI-PLC sample. Its localization to the surface along with having a PAN domain signifies that it is likely a receptor binding protein making it and the molting protein Bm3215 potential therapeutic or diagnostic targets.

The parasitic nematode-specific proteins highlight proteins involved in the ability of these parasites to successfully live and reproduce within their hosts and the filarial nematode specific proteins presumably have even more specialized roles. Bm13388 is interesting because, to date, the only homology for this protein is in *Strongyloides ratti*, *Loa loa* and *Onchocerca volvulus*. Bm13388 was only found in the membrane PI-PLC sample and is predicted to be GPI-AP by both GPI-SOM and bigPI. It has been shown to be one of the top fifteen proteins expressed in the extracellular vesicle proteome of adult females [53]. Since this protein is released into the host in extracellular vesicles and is specific to parasite nematodes, it may play an important role in the way these parasitic nematodes interact with the host.

We compared the 1012 proteins we compiled from all LC-MS/MS data to previously published proteomic studies and found new evidence for the expression of 84 *B*. *malayi* and 3 *Wolbachia* proteins. These are noted in S7 Table with the Wormbase ID of the novel identified proteins highlighted in red for 29 proteins when a synonym ID for comparison was found or in green for 58 proteins when no synonym was found, likely signifying a new gene model in updated genome versions. For the 954 proteins that had matches with previous proteomes, the intersection data is summarized with UpSetR [50] analysis in S3 Fig. As expected from our similar analysis with the predicted GPI-APome (Fig 3B), there were fewer matches with proteins from excretory-secretory proteomes [54, 55] and the extracellular vesicle proteome [53]. The majority of the matches (944 of 954) were found in the two largest proteome datasets: the 2011 stage specific proteome set [57] and the 2015 proteomes of the body wall, digestive tract and reproductive tract [56]. Thus, the corroborating proteins we identified in these previously published proteomes gives additional insights on their life stage and/or tissue expression.

The *C. elegans* lipid raft study identified 44 proteins of which 21 were predicted to be GPI-APs [61]. We found *B. malayi* orthologs for 17 of these 44 lipid raft associated proteins in our LC-MS/MS data (S7 Table) showing that our method also captured lipid raft proteins. Of these 17 proteins, 4 were predicted to be GPI-APs in *C. elegans* [61] while two of the *B. malayi* orthologs (Bm2418 and Bm8928) were predicted GPI-APs in our computational GPI-APome. Similarly, in our LC-MS/MS data, we found *B. malayi* orthologs for 7 out of 22 proteins identified in a *C. elegans* GPI-AP proteomic study [17]. Of these seven, four (Bm1904, Bm1972, Bm2418, Bm5444) were present in our predicted *B. malayi* GPI-APome. This variability in GPI-AP prediction suggests that the prediction programs give a reasonable first indication of potential GPI-APs. Combined with experimental approaches, such as those reported here, a more comprehensive representation of the GPI-APome is gained.

## Conclusions

Our study both computationally and experimentally defines the GPI-APome in the parasitic nematode *B. malayi,* the first such study in any parasitic worm. This proteome includes both novel and previously identified GPI-anchored proteins that likely play an important role during the parasite’s lifecycle. Better understanding of the GPI-APome and its synthesis pathway may lead to the identification of new strategies for diagnosis and treatment for this important parasite.

## Acknowledgements

We thank Dr. Zhiru Li for *E. coli* OP50 and *C. elegans* N2, Dr. Saulius Vainauskas for PI-PLC enzyme, and Dr. Saulius Vainauskas, Dr. Clotilde S. Carlow, Dr. William Jack and Dr. Thomas C. Evans Jr. for critical feedback and proofreading of the manuscript. We thank Dr. Donald Comb and Mr. James Ellard for continued support of parasite glycobiology research.

## Supporting information

**S1 Fig. Dolichol phosphate mannosyltransferase comparison.** The DPM proteins for human, *S. cerevisiae, T. brucei, C. elegans* and *B. malayi* are shown with predicted transmembrane regions highlighted in yellow. Protein ID and protein length (aa = amino acids) are indicated for each.

**S2 Fig. Mock Control Dot Blot.** *B. malayi and C. elegans* lysates show no visible background signal or autofluorescence when FLAER reagent is not present. Samples: BSA, *E.coli* OP50 total and soluble lysate, *C. elegans* N2 total and soluble lysate, *B. malayi* total lysate 1: B. *malayi* total lysate mock control, *B. malayi* total lysate 2: *B. malayi* surface PI-PLC treated total lysate. After 1:1 series dilution, samples were spotted on to nitrocellulose membrane and incubated overnight in buffer.

**S3 Fig. UpSetR Proteomic Comparison of LC-MS/MS data to published proteomics data.** EV 2018 [53] is an extracellular vesicle proteome. ES 2008 [54] and ES 2009 [55] are excretory and secretory proteomes and BDR 2015 [56] is body wall, digestive tract and reproductive tract proteomes. SS 2011 [57] is a stage specific proteome set. Proteins that are exclusive to a set are designated with a single dot. Proteins that are found in multiple sets have dots that are linked with lines. The set size shown on the left is the number of proteins identified in the different proteome datasets that match the 1012 proteins identified in this study.

**S1 Table. *B. malayi* predicted GPI-APome.** List of proteins predicted to be *B. malayi* GPI-APs along with the prediction program(s) that identified that protein. Proteins that did not have N-terminal sequence or had more than three predicted transmembrane domains were removed. Wormbase ID is used as an identifier for each protein. C-terminal cleavage site / omega site predictions and quality scores are included when information was available. We compiled the available synonym identifiers and utilized those identifiers to compare the presence of these predicted GPI-APs with previously published *B. malayi* proteomes. The proteins that were identified in our LC-MS/MS data are highlighted in yellow.

**S2 Table. *B. malayi* Surface Mock Control LC-MS/MS.** List of proteins identified from adult *B. malayi* not treated with PI-PLC enzyme. Proteins from the intact worms were identified using LC-MS/MS. Spectral data was searched against *B. malayi* and its endosymbiont Wolbachia using Byonic software from Protein Metrics to identify proteins. Rows highlighted in yellow match a protein identified in *B. malayi* predicted GPI-APome. Proteins that only had one unique peptide are highlighted in red and were not used in further analysis.

**S3 Table. *B. malayi* Surface PI-PLC LC-MS/MS.** List of proteins identified from adult *B. malayi* treated with PI-PLC. Proteins released from the intact worms were identified using LC-MS/MS. Spectral data was searched against *B. malayi* and its endosymbiont Wolbachia using Byonic software from Protein Metrics to identify proteins. Rows highlighted in yellow match a protein identified in *B. malayi* predicted GPI-APome. Proteins that only had one unique peptide are highlighted in red and were not used in further analysis.

**S4 Table. *B. malayi* Membrane Mock control LC-MS/MS**. List of proteins identified from adult *B. malayi* membrane extracts not treated with PI-PLC enzyme. Proteins released from the membrane extracts were identified using LC-MS/MS. Spectral data was searched against *B. malayi* and its endosymbiont Wolbachia using Byonic software from Protein Metrics to identify proteins. Rows highlighted in yellow match a protein identified in *B. malayi* predicted GPI-APome. Proteins that only had one unique peptide are highlighted in red and were not used in further analysis.

**S5 Table. *B. malayi* Membrane PI-PLC LC-MS/MS**. List of proteins identified from adult *B. malayi* membrane extracts treated with PI-PLC. Proteins released from the membrane extracts were identified using LC-MS/MS. Spectral data was searched against *B. malayi* and its endosymbiont Wolbachia using Byonic software from Protein Metrics to identify proteins. Rows highlighted in yellow match a protein identified in *B. malayi* predicted GPI-APome. Proteins that only had one unique peptide are highlighted in red and were not used in further analysis.

**S6 Table. *B. malayi* Membrane Enrich GPI-AP LC-MS/MS**. List of proteins identified from adult *B. malayi* membrane extracts treated with a series of organic solvents to isolate GPI-AP enriched fractions. Enriched proteins were identified using LC-MS/MS. Spectral data was searched against *B. malayi* and its endosymbiont Wolbachia using Byonic software from Protein Metrics to identify proteins. Rows highlighted in yellow match a protein identified in *B. malayi* predicted GPI-APome. Proteins that only had one unique peptide are highlighted in red and were not used in further analysis.

**S7 Table. *B. malayi* compiled LC-MS/MS with proteomic comparison**. Proteins identified from the *B. malayi* surface and membrane samples treated with or without PI-PLC and membrane sample enriched for GPI-AP using organic solvents were compiled into a single list if the protein had two or more unique peptides. Compiled available synonym identifiers were utilized to compare the presence of these proteins with previously published *B. malayi* proteomes. The Wormbase ID of the proteins not present in previously published *B. malayi* proteomes are highlighted in red when a synonym ID for comparison was found or in green when no synonym was found. Rows highlighted in yellow match a protein identified in *B. malayi* predicted GPI-APome. In addition, we noted if the protein was an ortholog to a protein identified in the *C. elegans* lipid raft proteome [61] or *C. elegans* GPI-AP study [17]. Proteins that match computationally predicted GPI-APome proteins are highlighted in yellow.

